## Supplementary figures and images for "A timer gene network is spatially regulated by the terminal system in the *Drosophila* embryo"

### A1-1.tiff

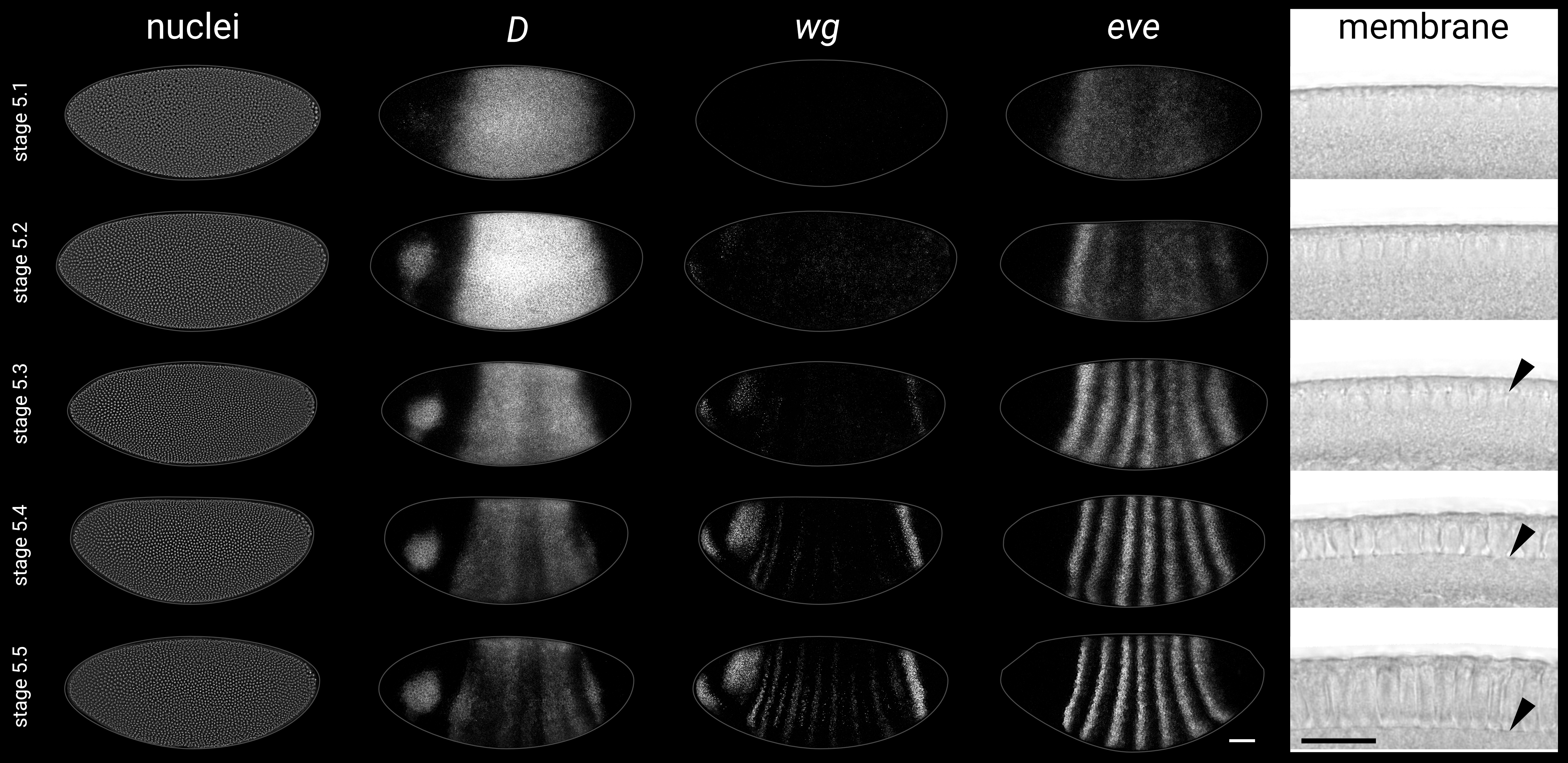

### A2-1.tiff

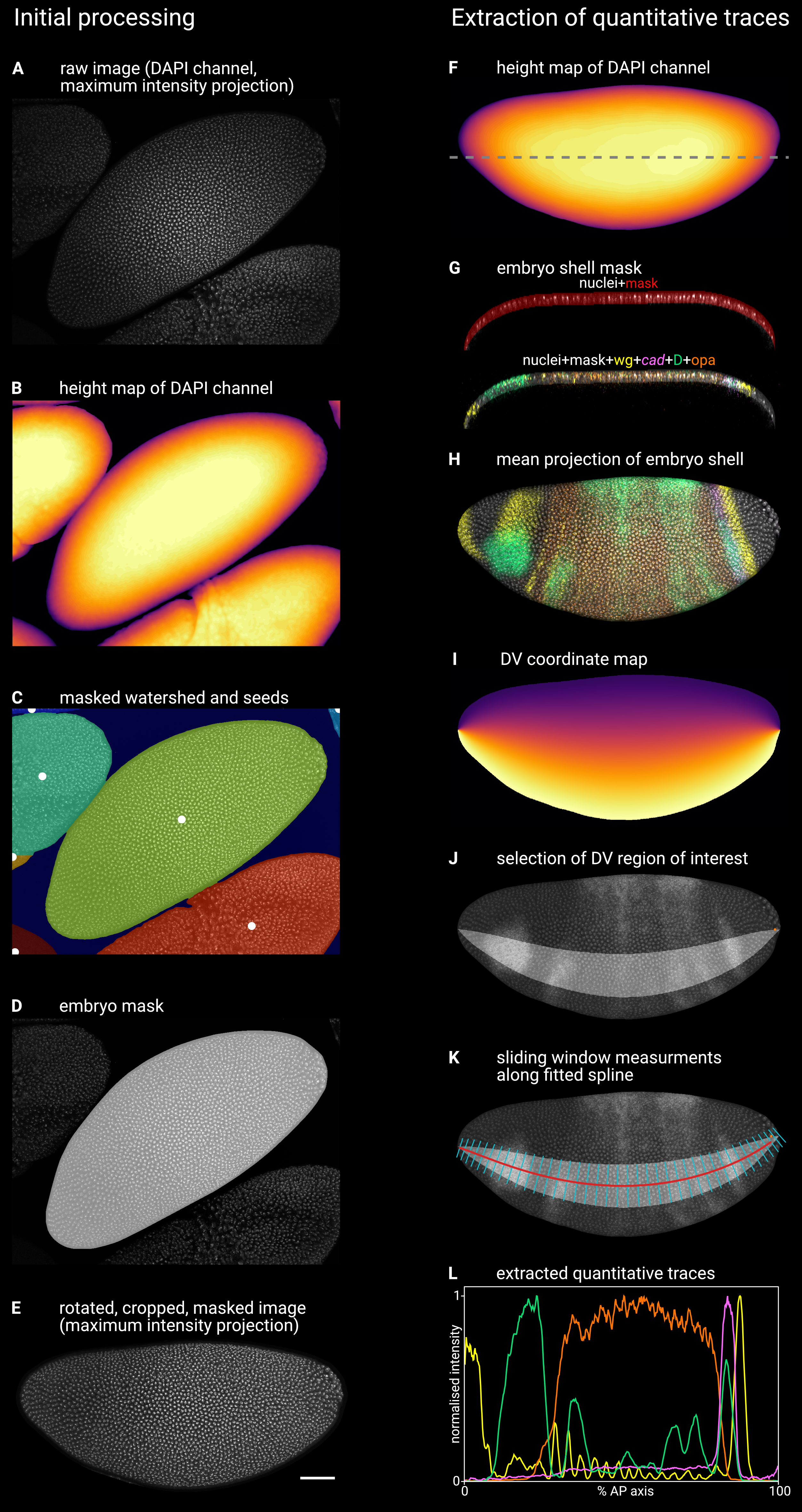

### s1-1.tiff

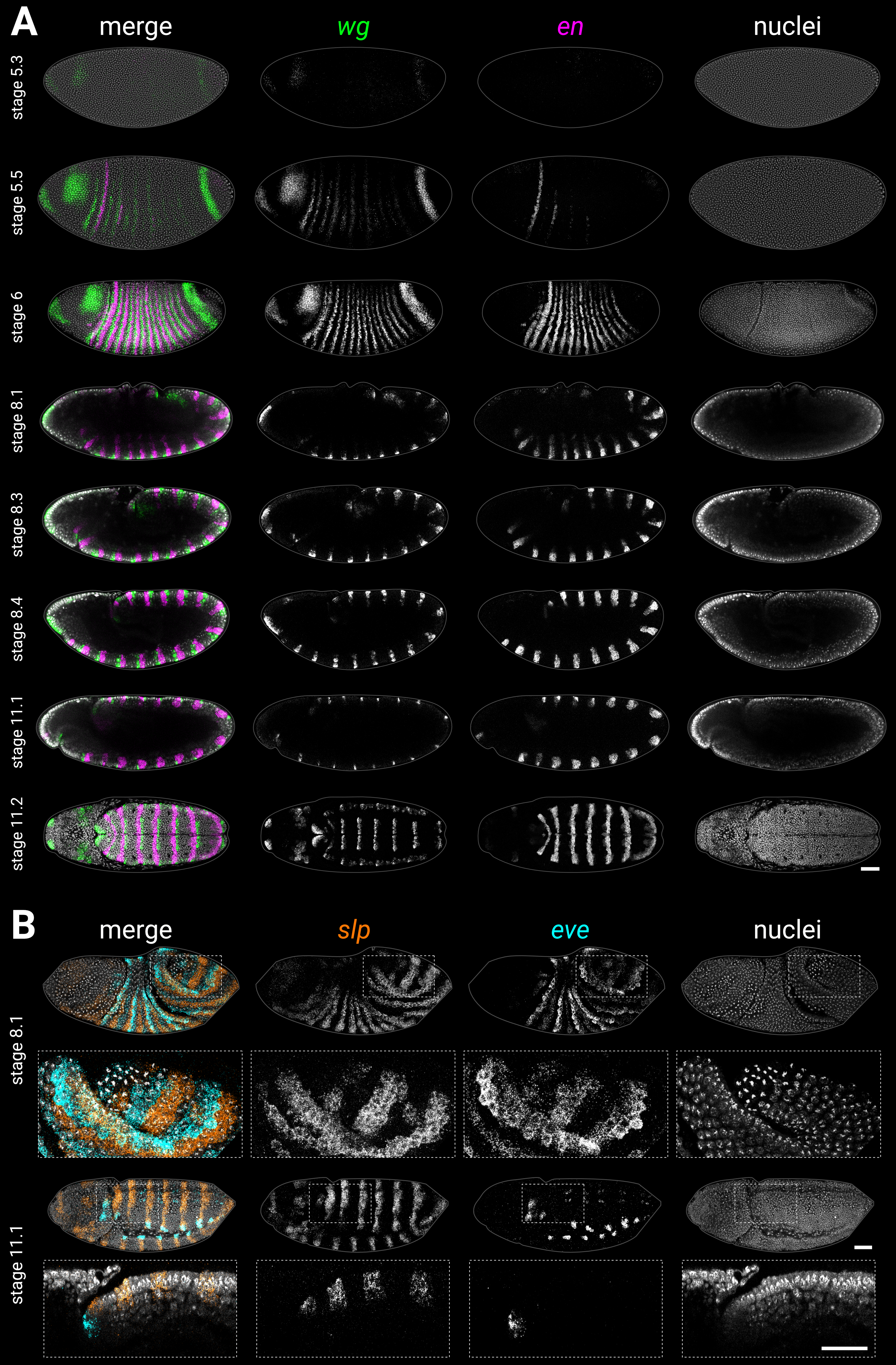

### s2-1.tiff

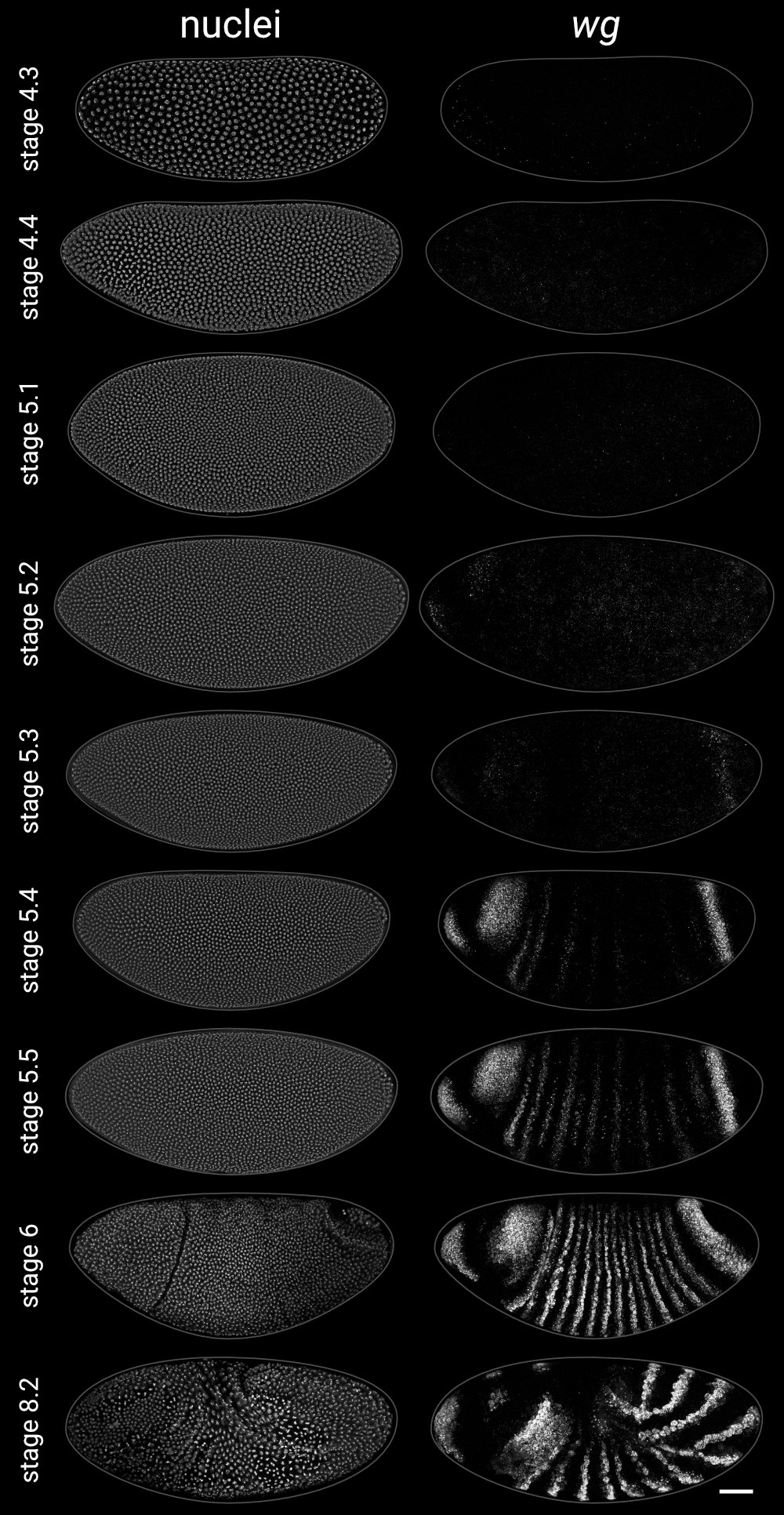

### s2-2.tiff

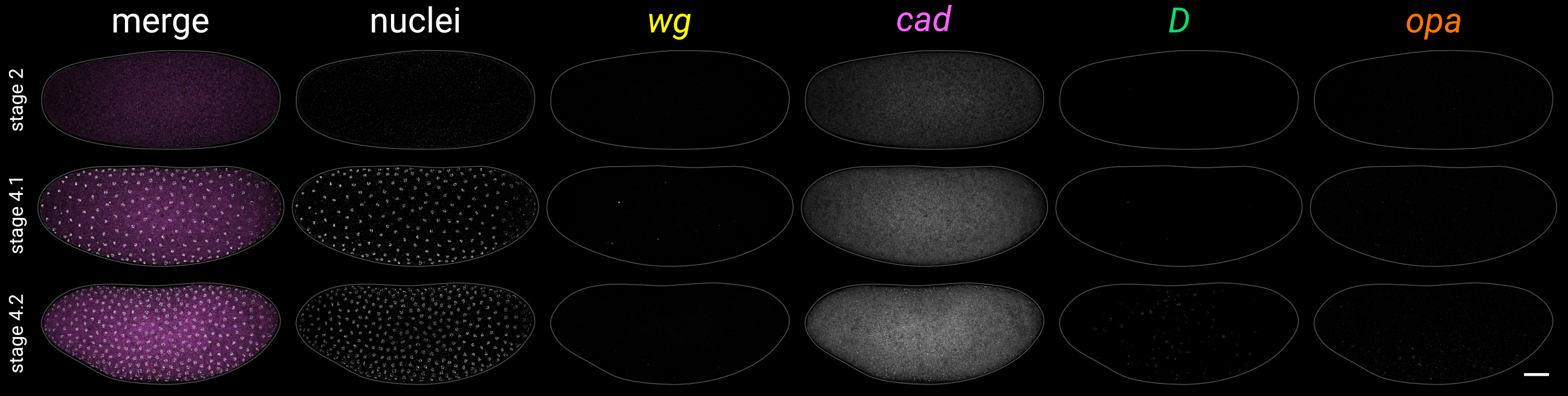

### s2-3.tiff

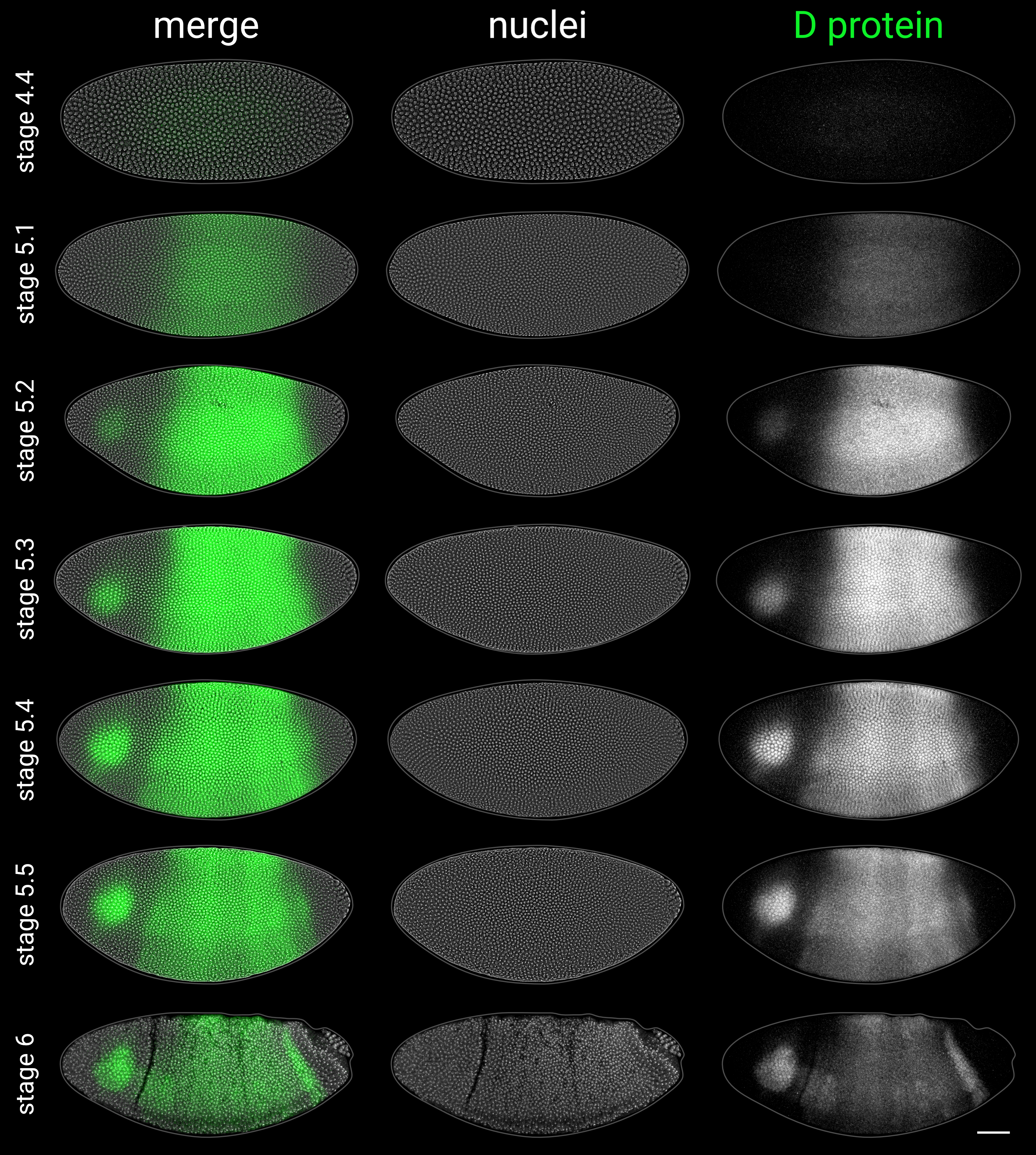

### s2-4.tiff

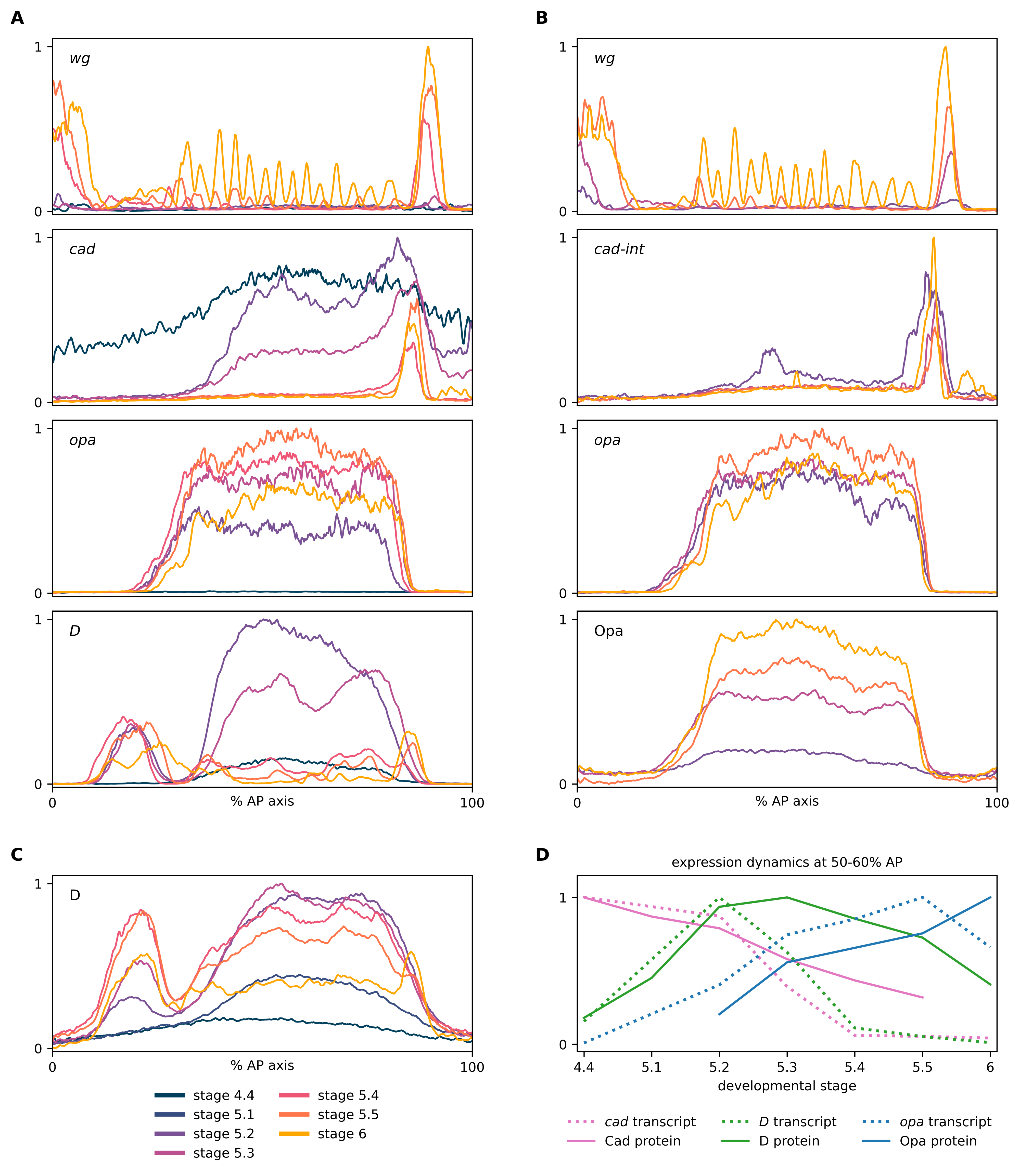

### s3-1.tiff

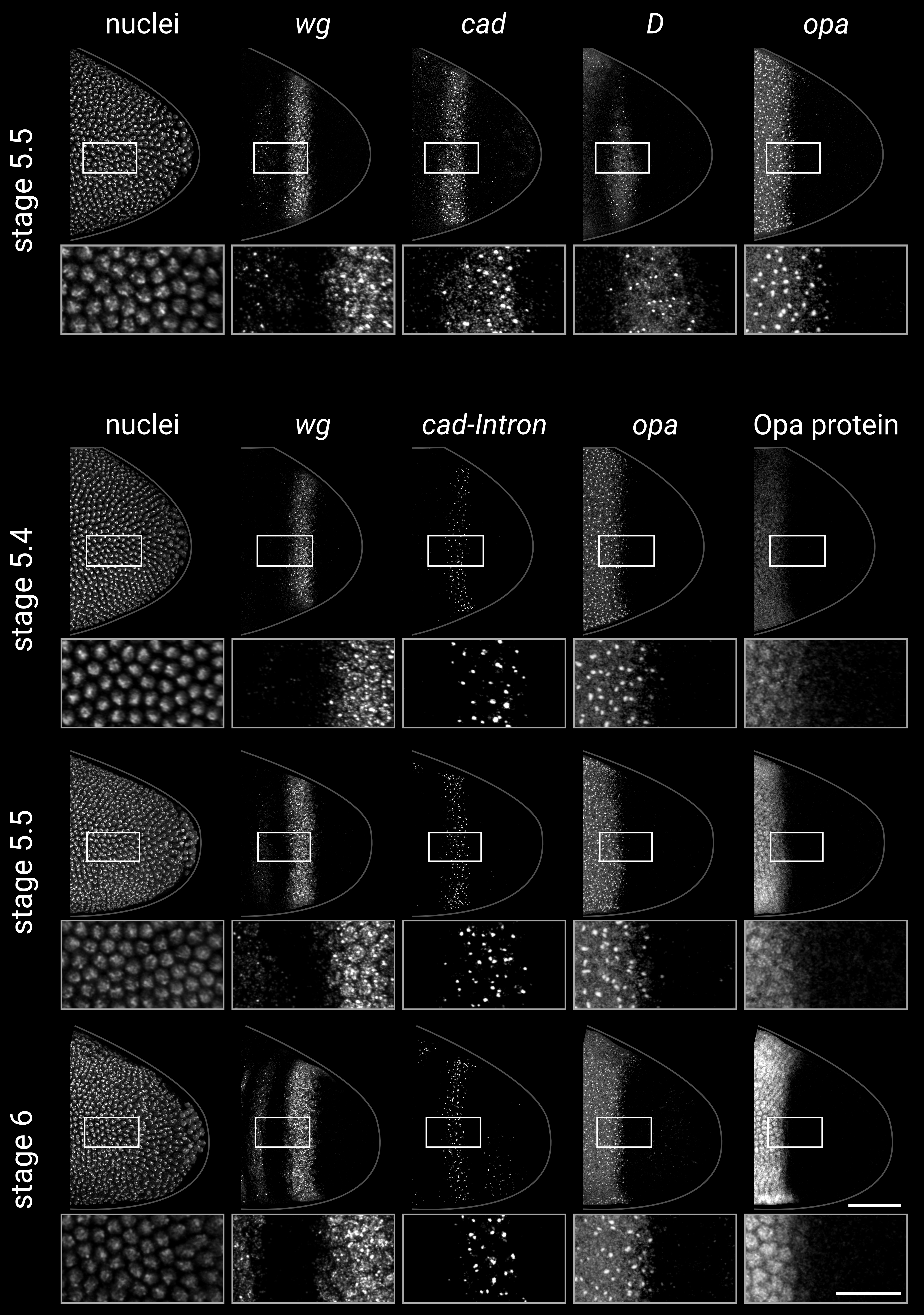

### s4-1.tiff

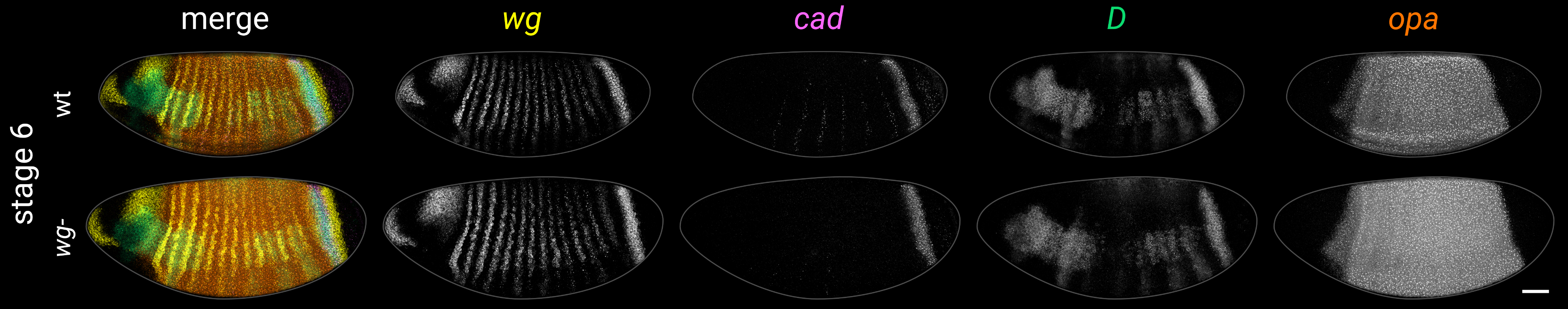

### s4-2.tiff

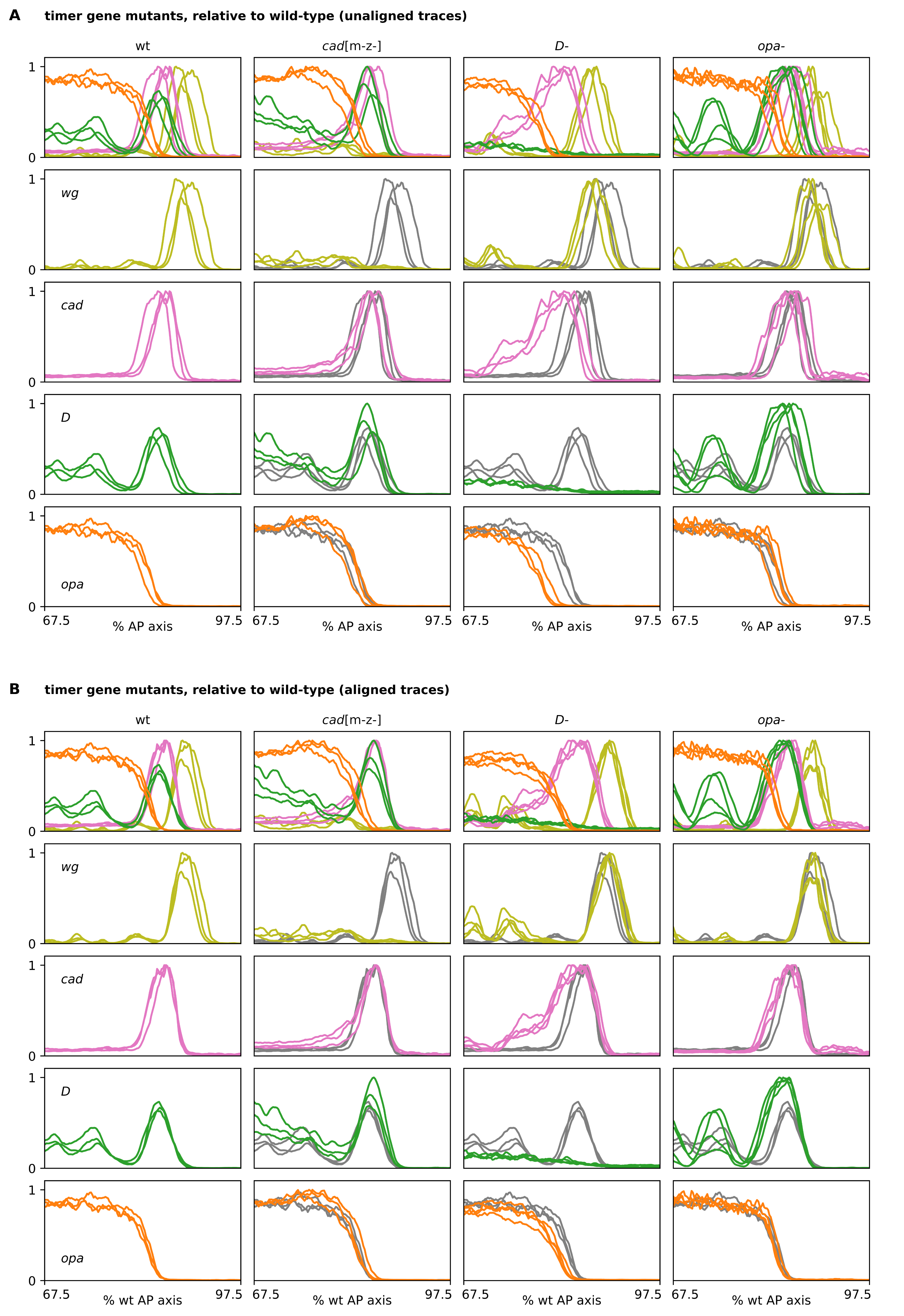

### s4-3.tiff

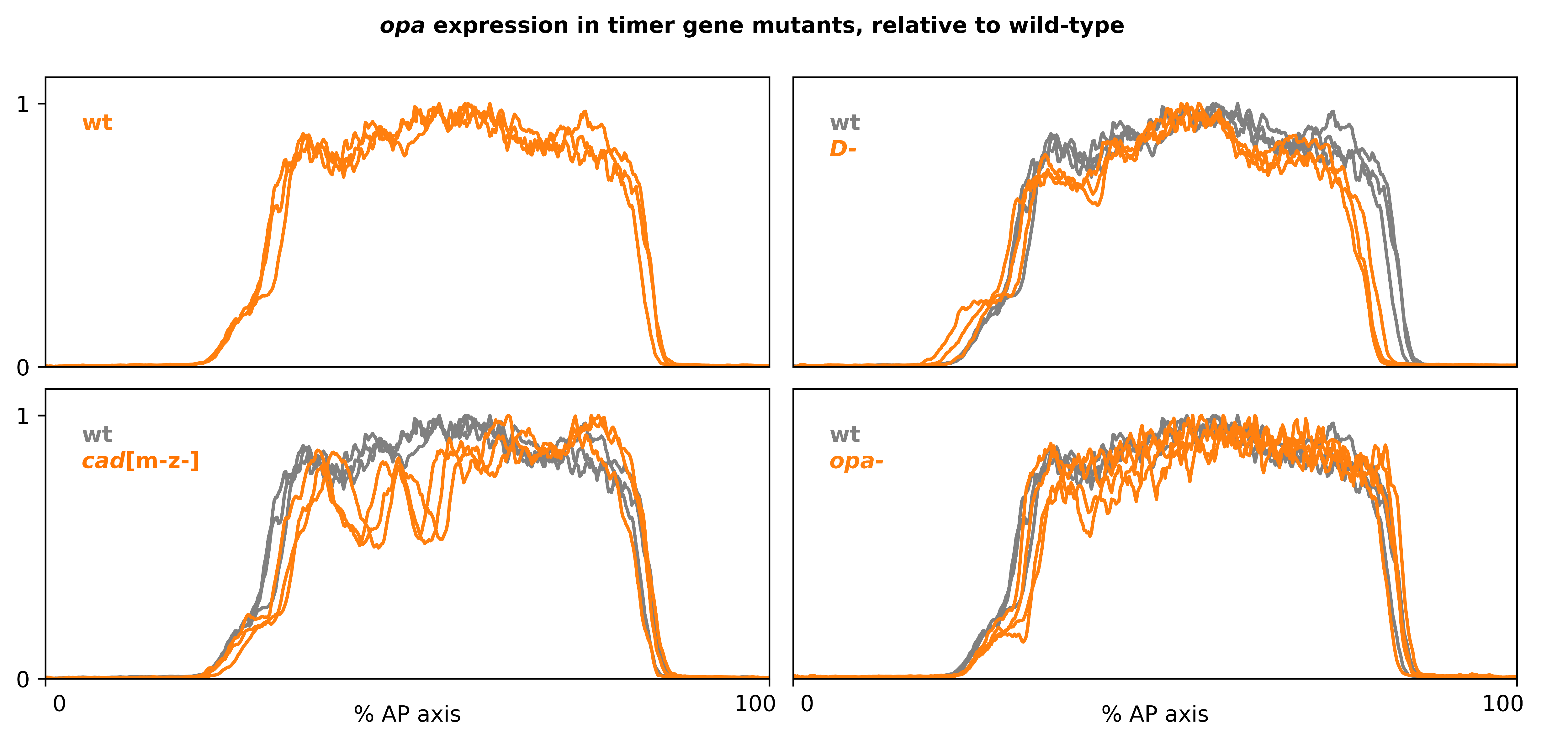

### s4-4.tiff

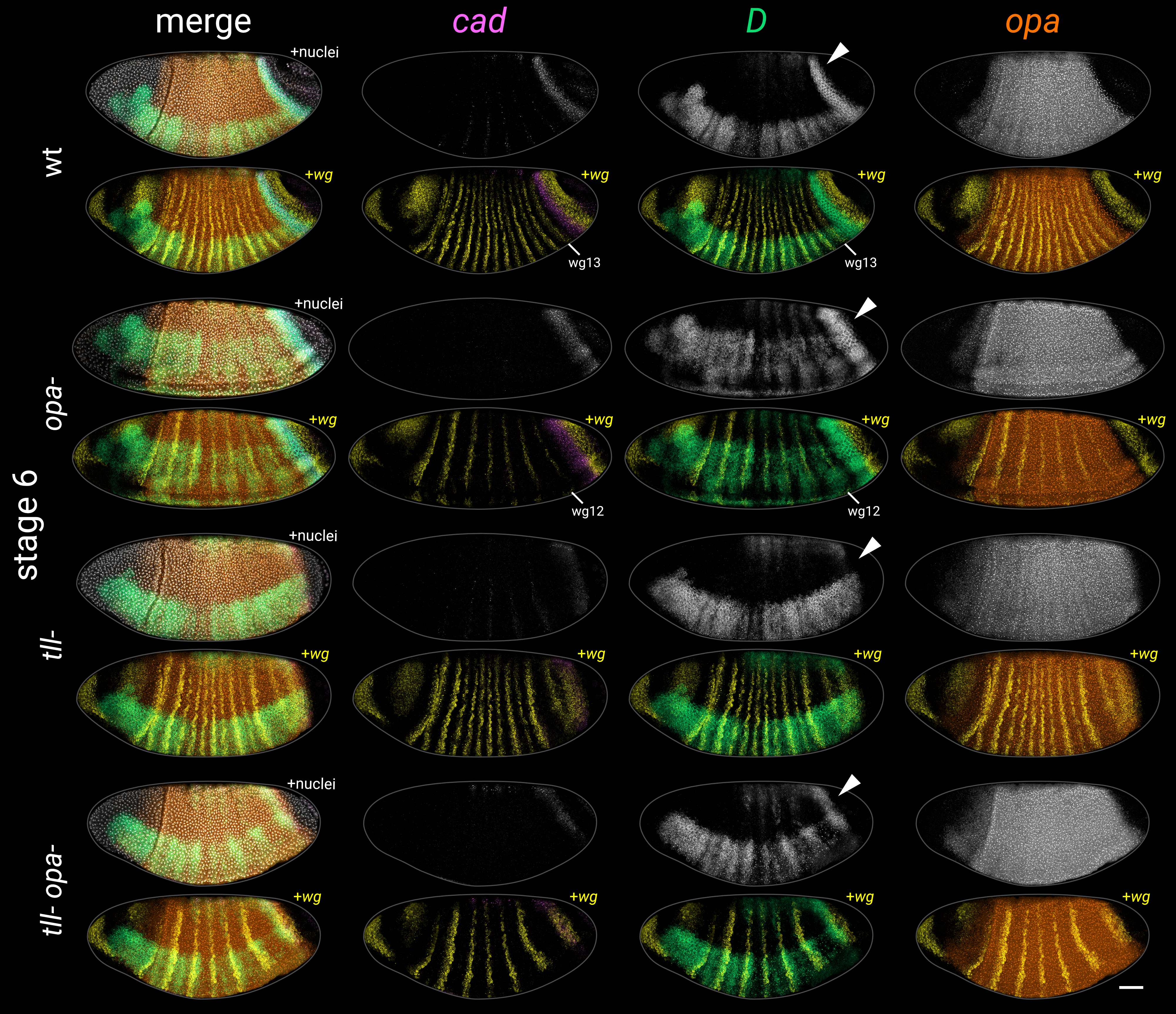

### s4-5.tiff

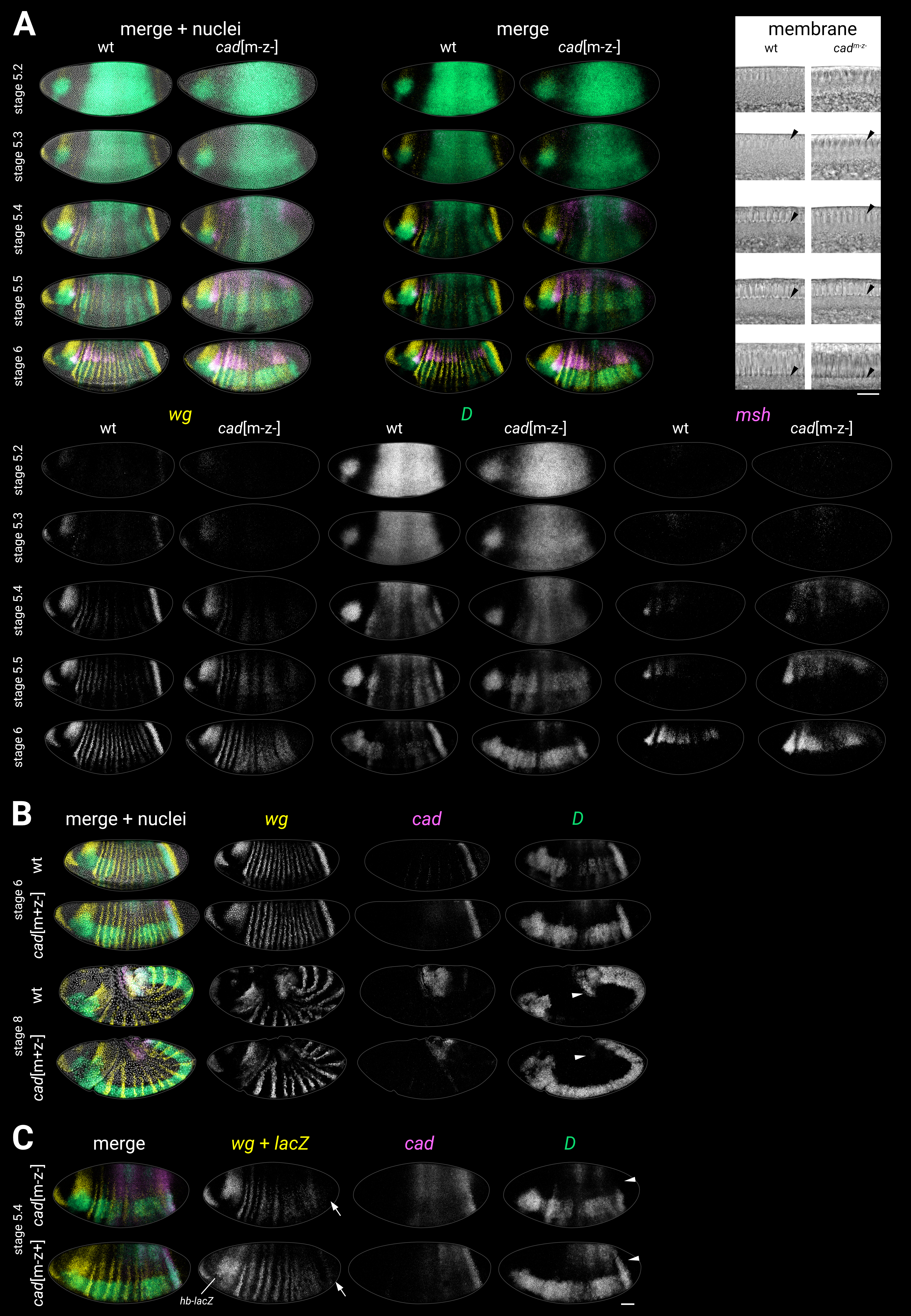

### s4-6.tiff

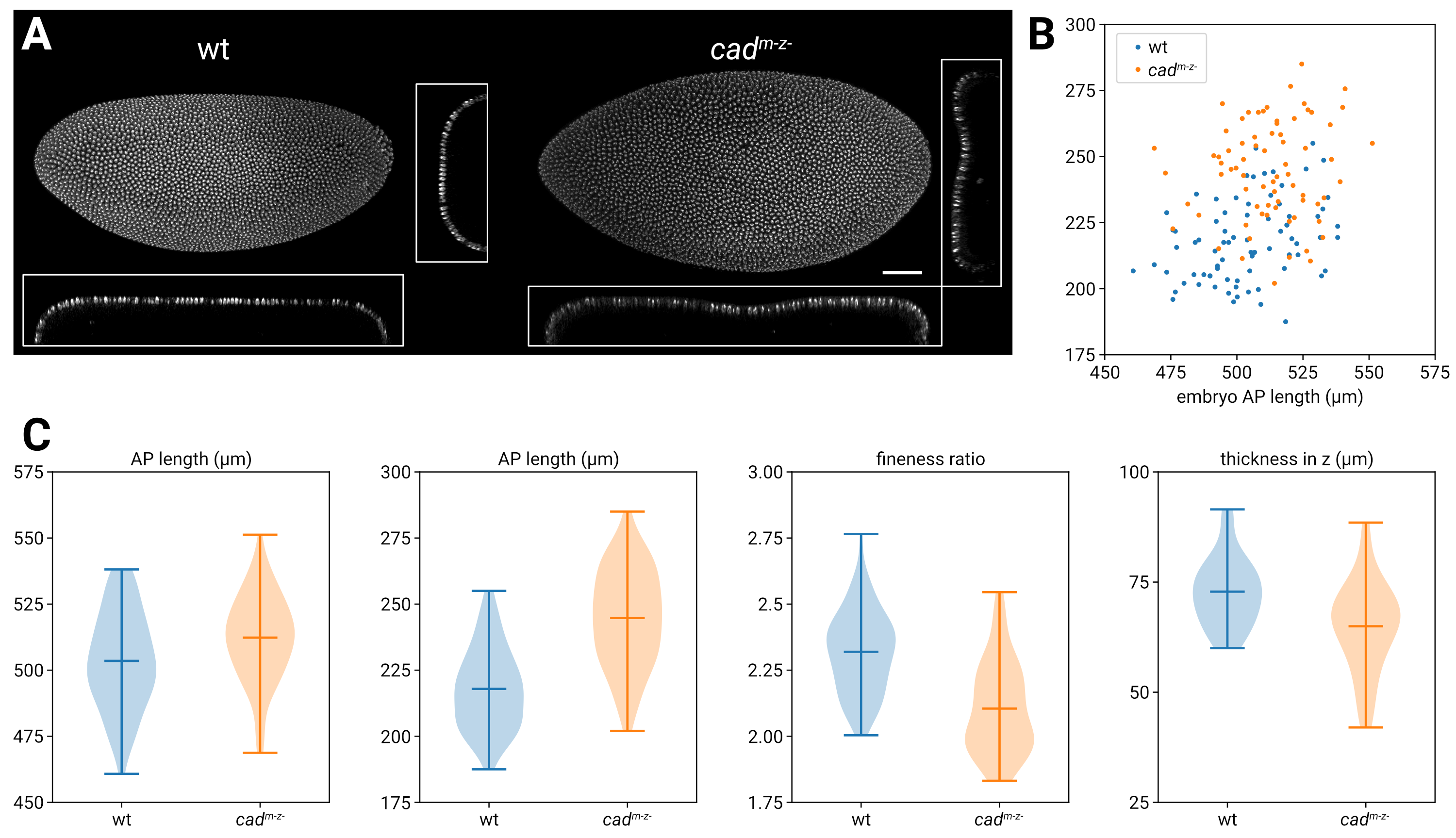

### s5-1.tiff

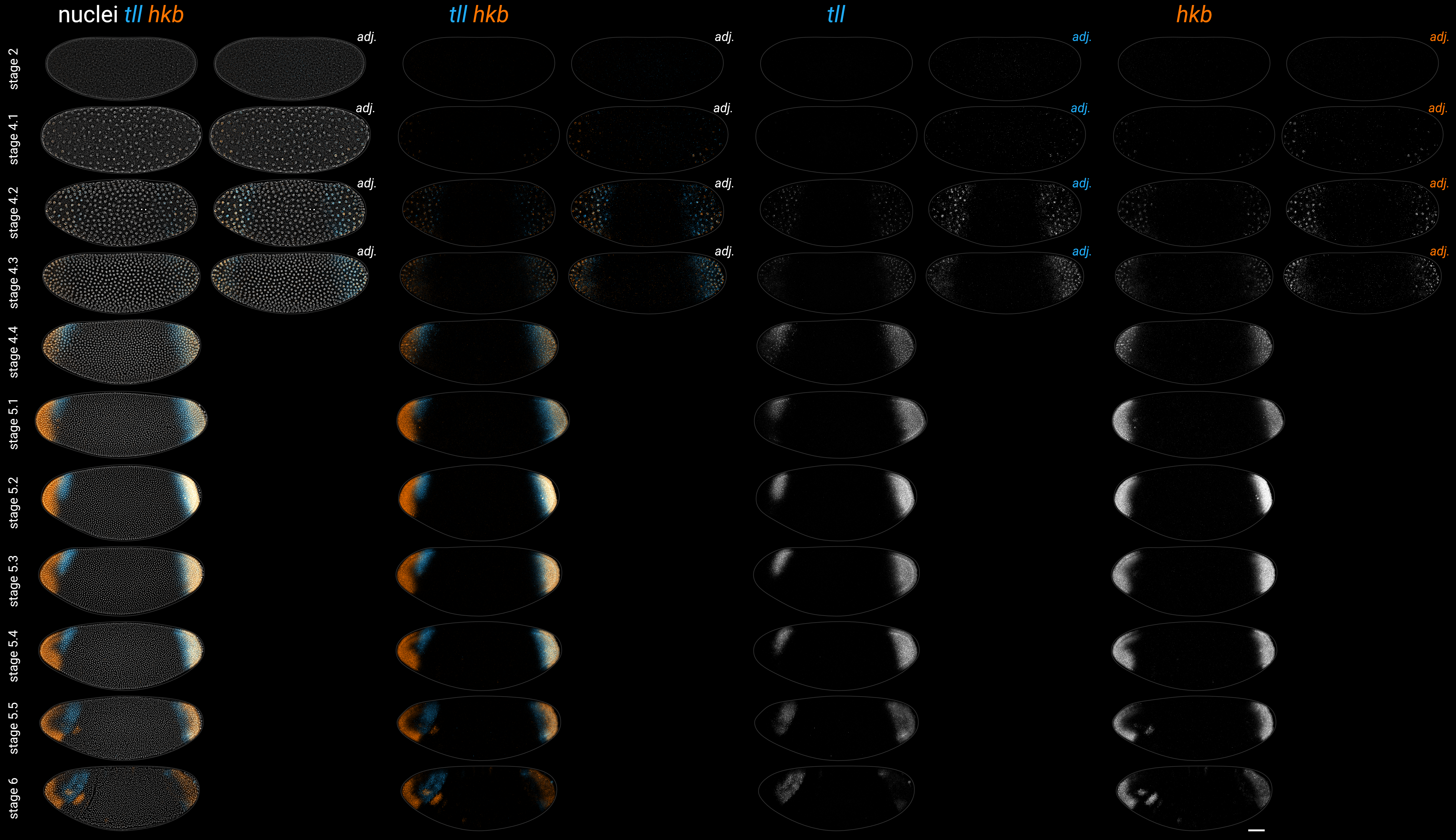

### s5-2.tiff

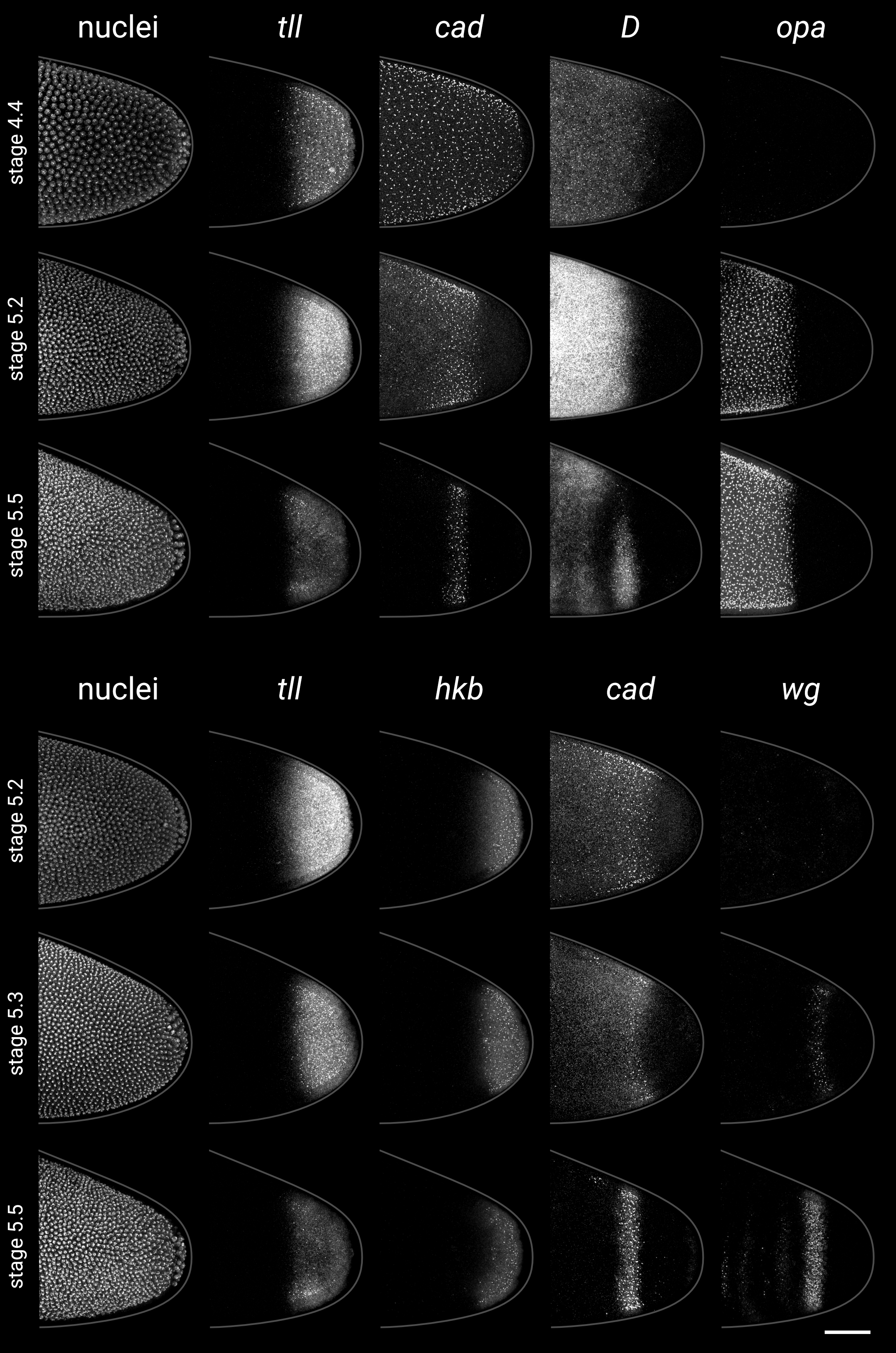

### s5-3.tiff

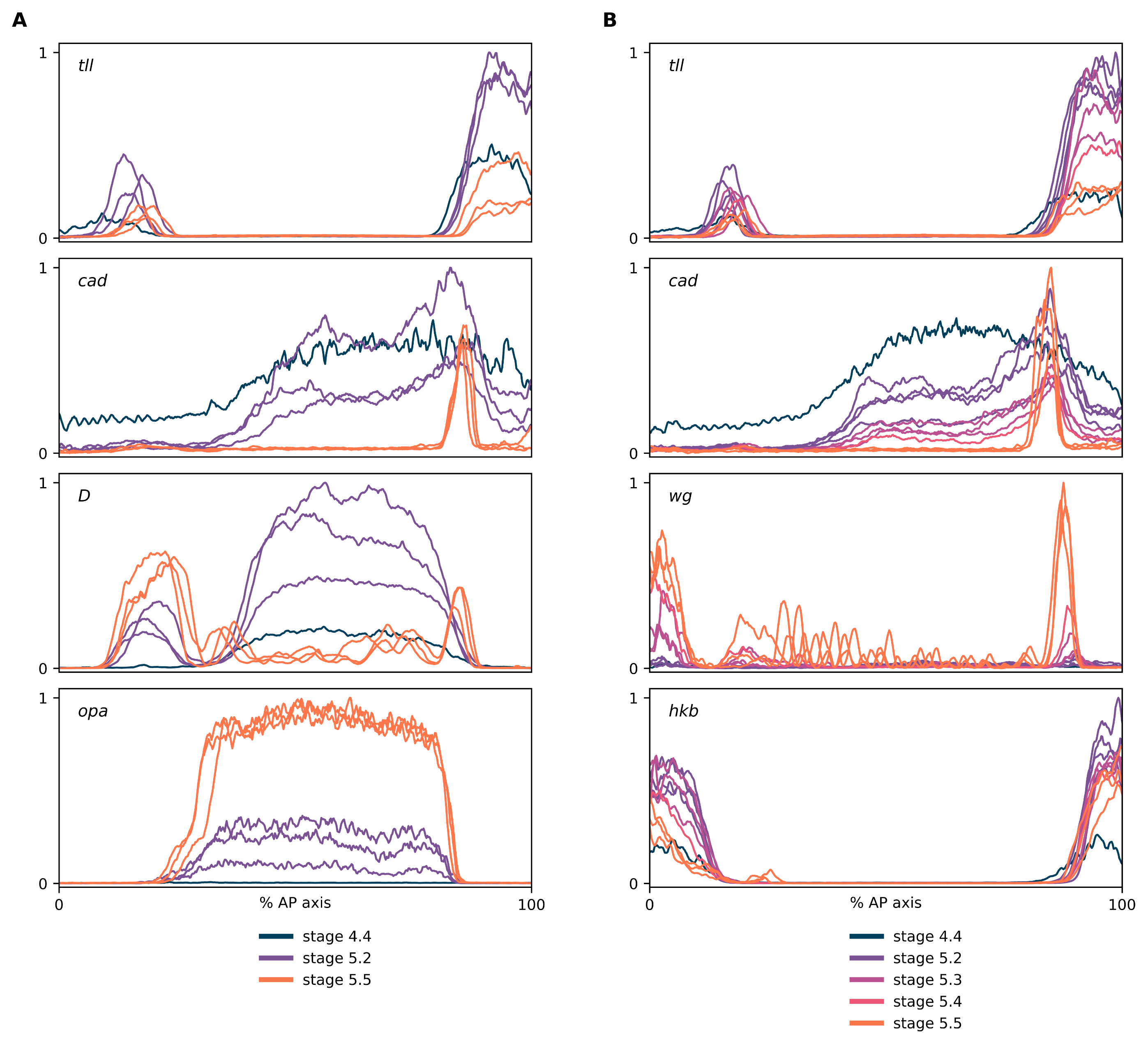

### s5-4.tiff

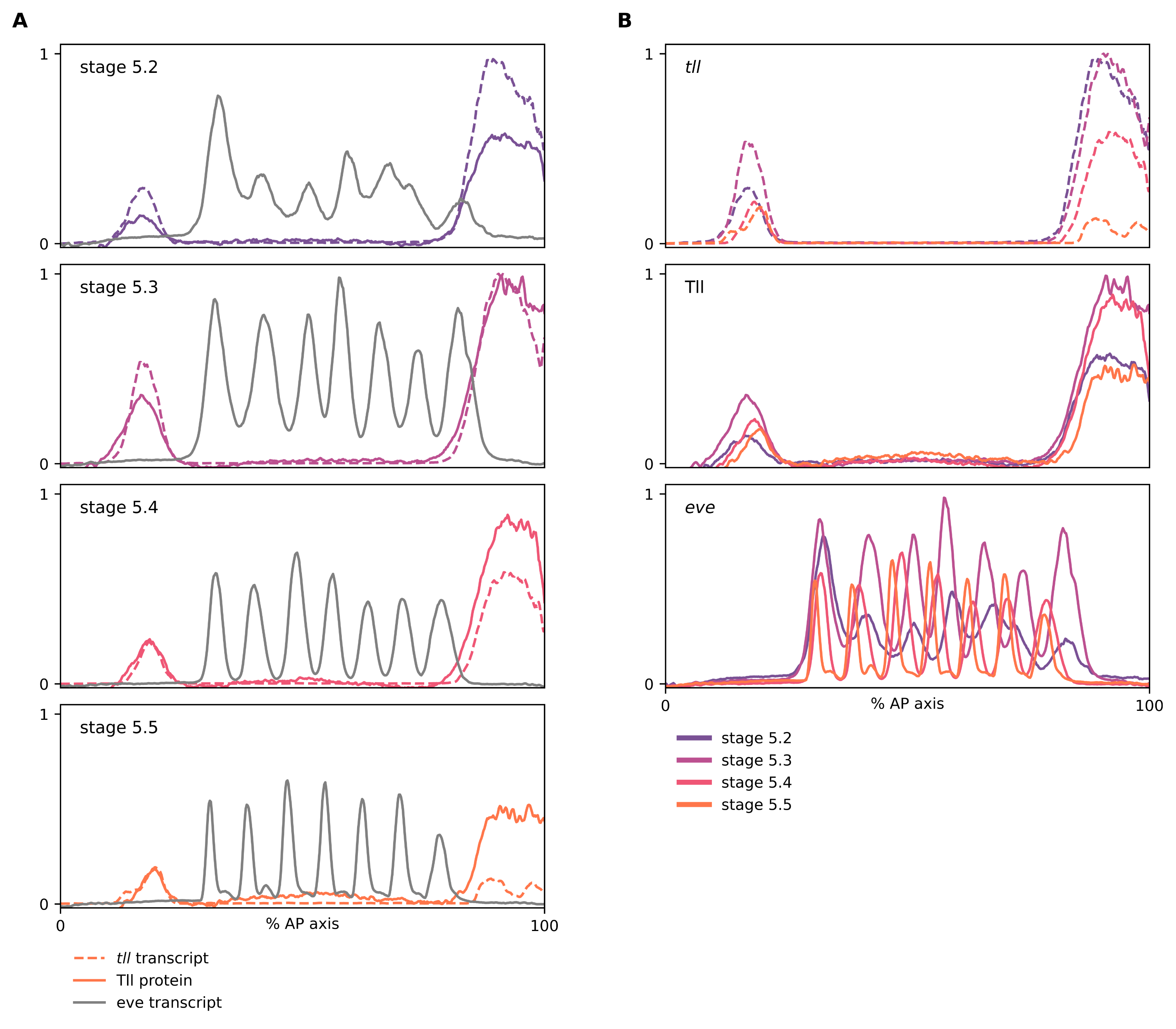

### s5-5.tiff

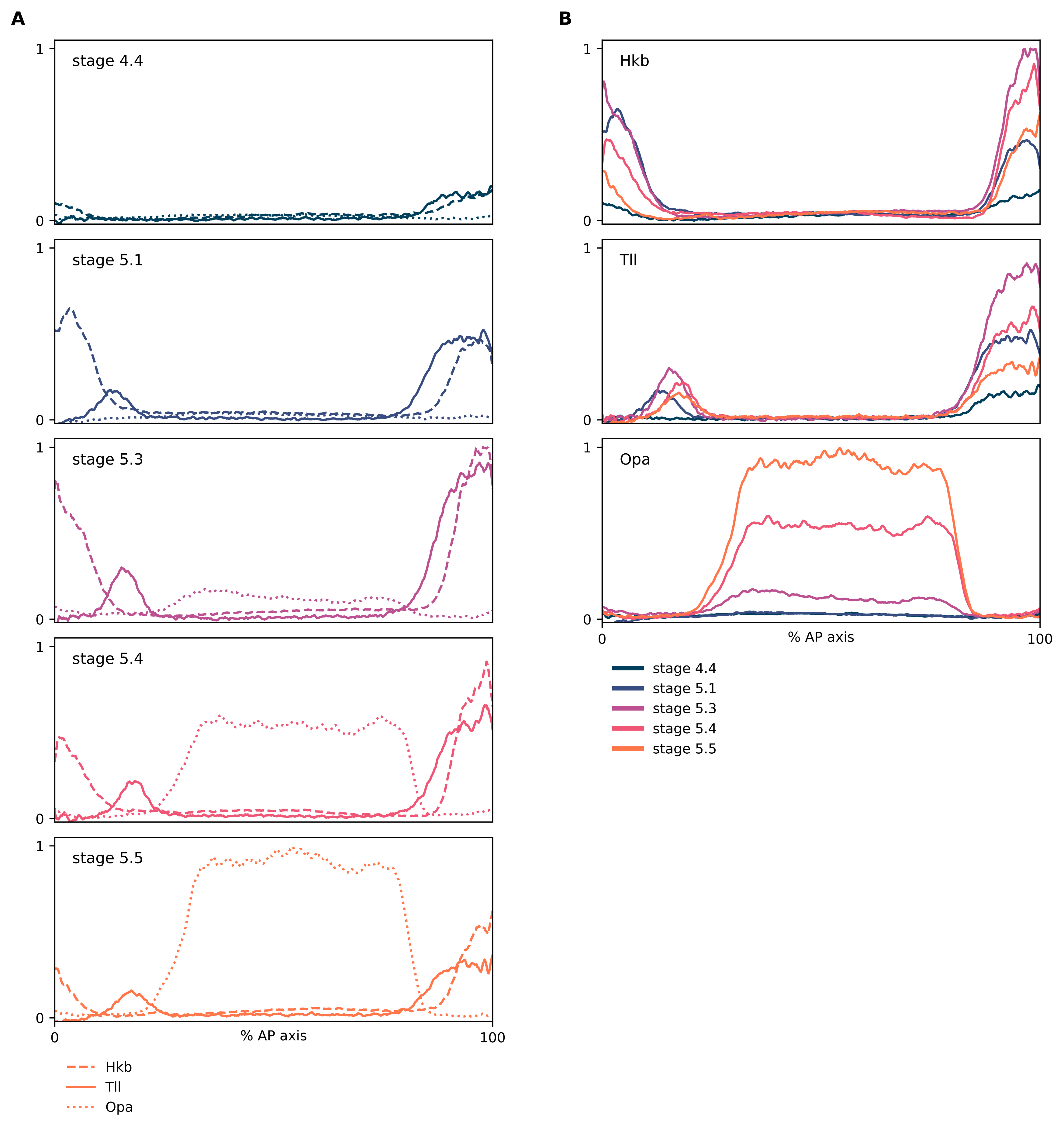

### s5-6.tiff

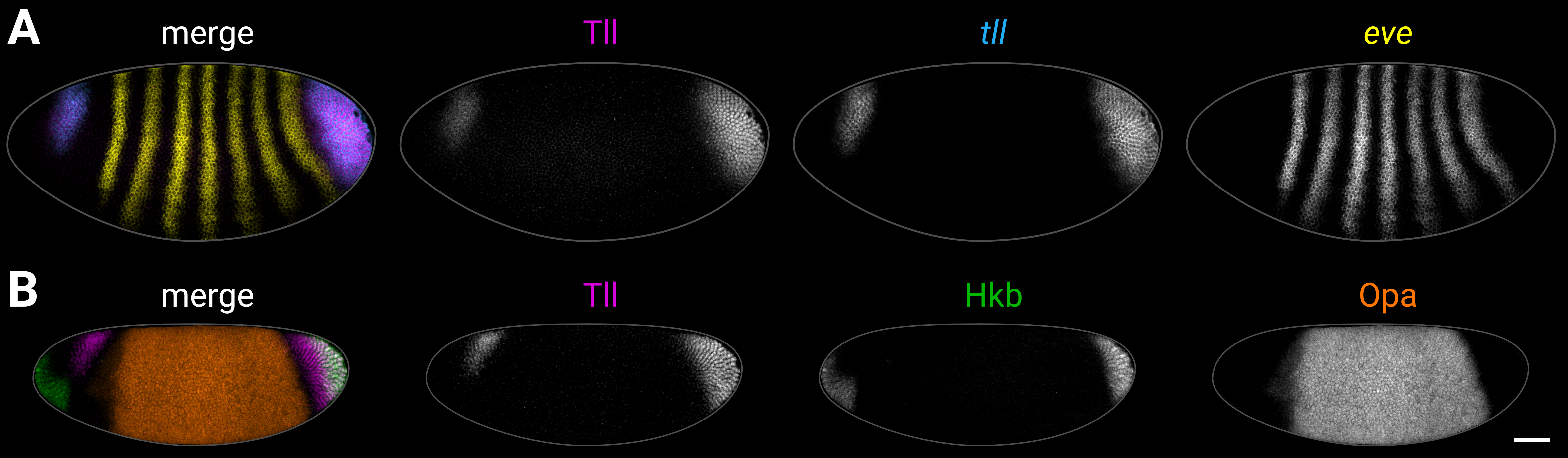

### s6-1.tiff

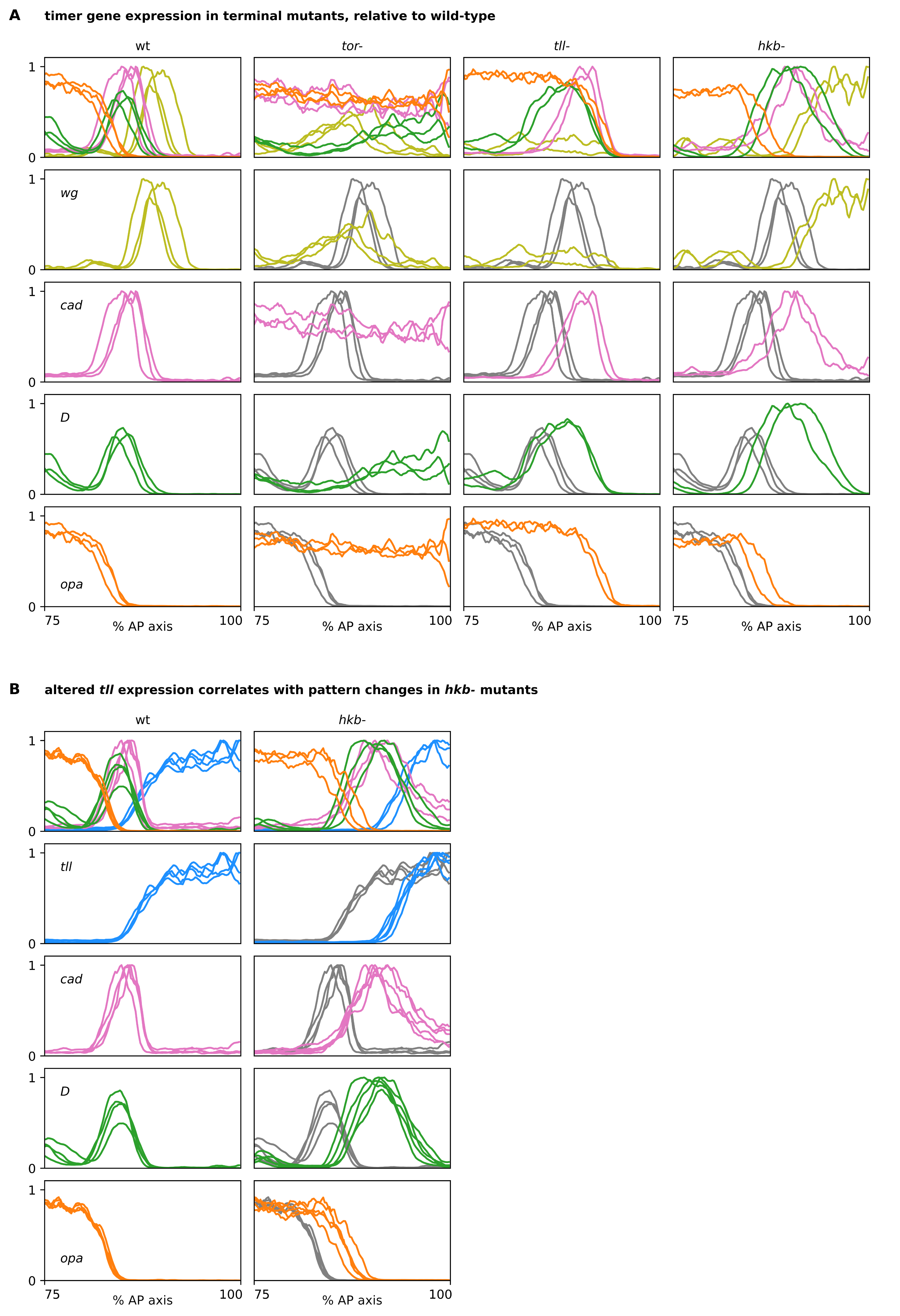

### s6-2.tiff

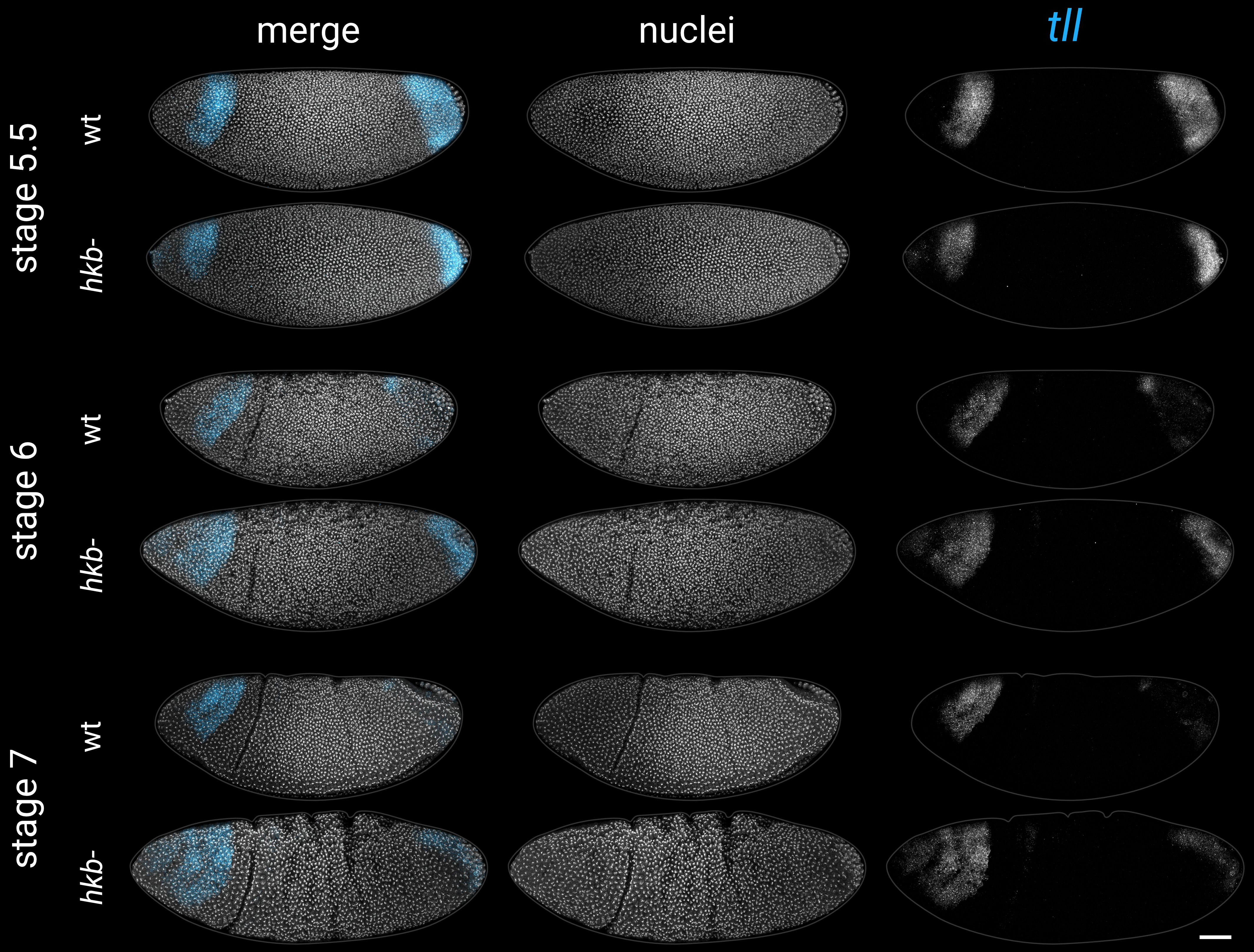

### s7-1.tiff

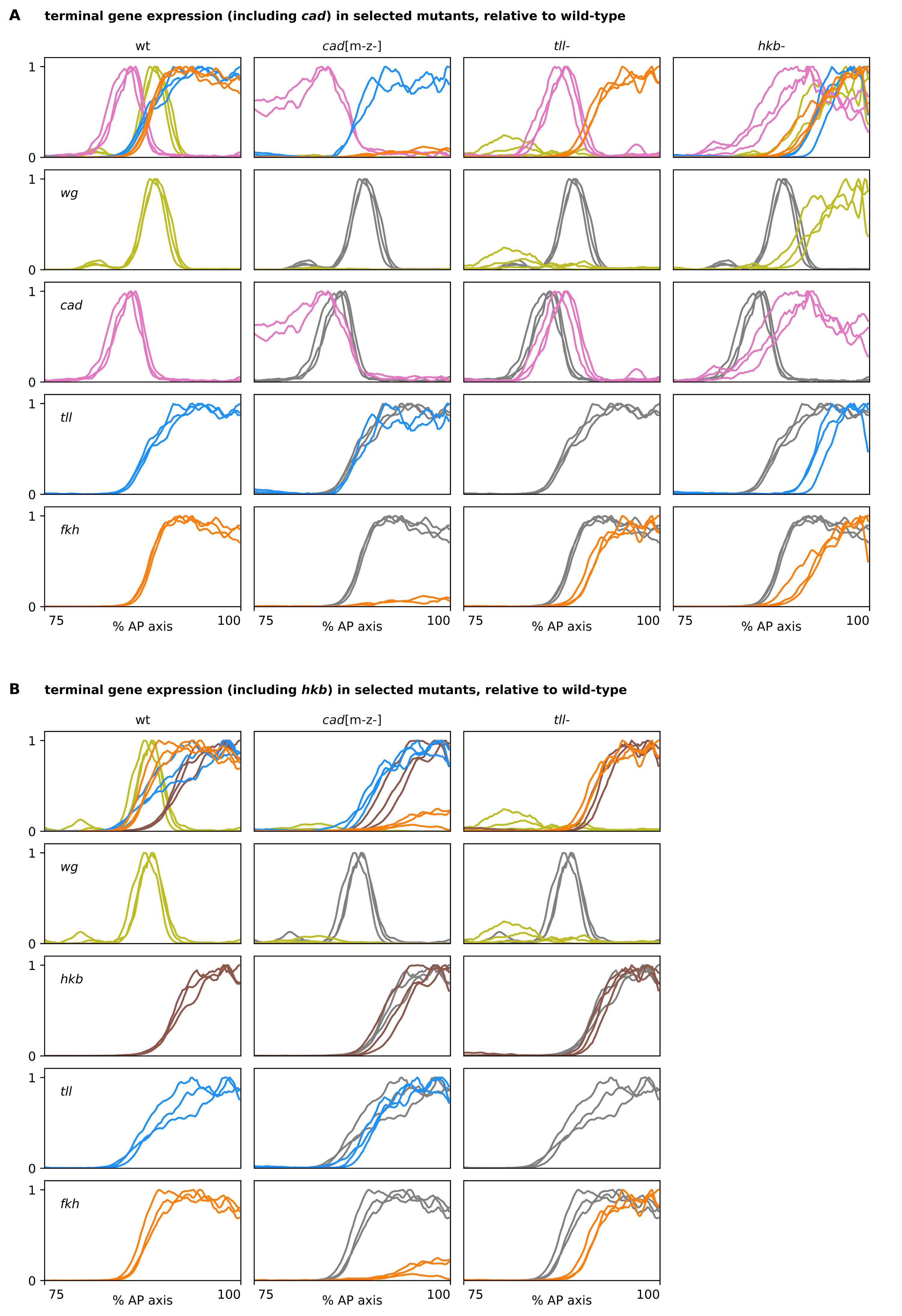

### s7-2.tiff

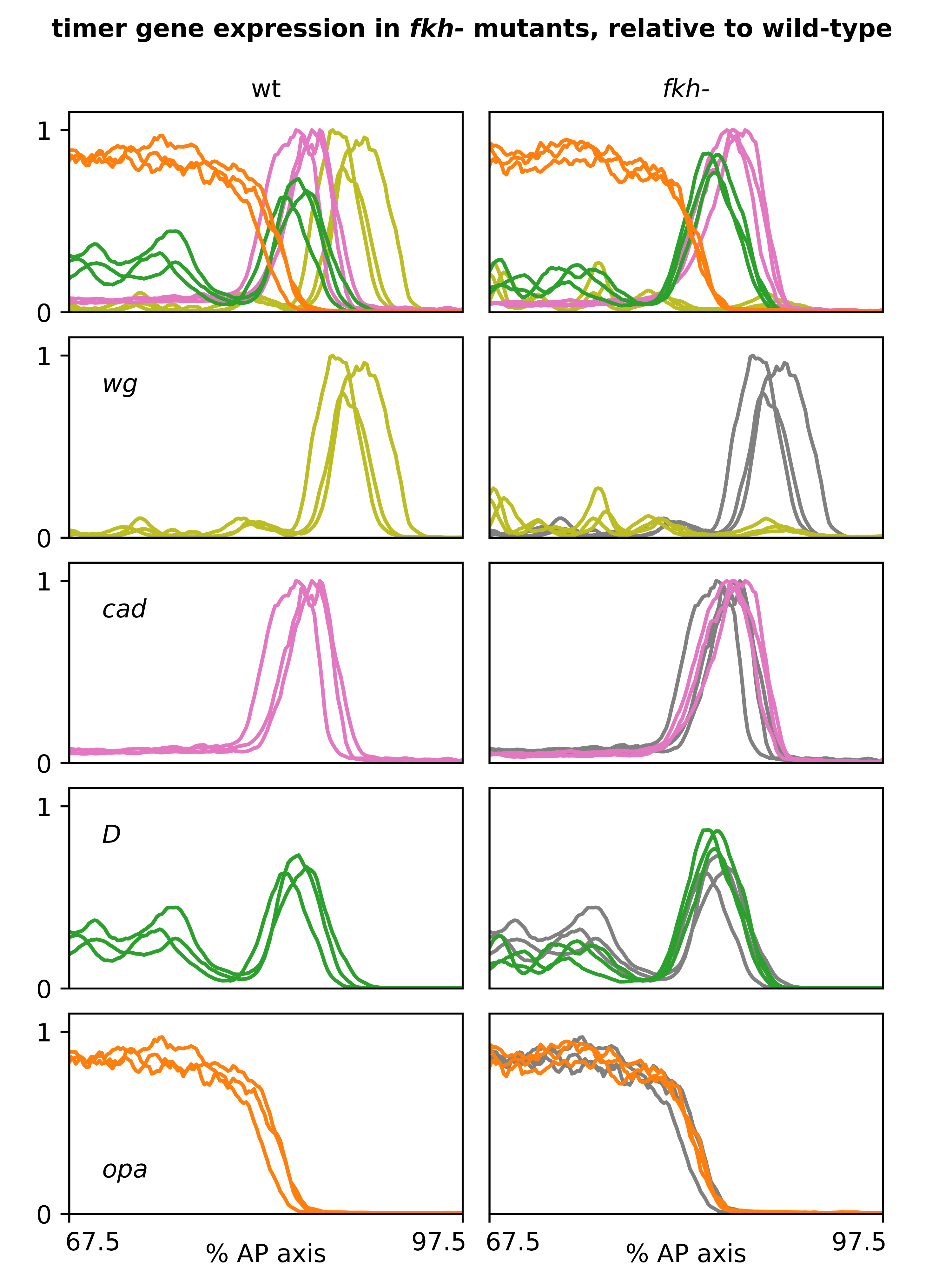

### s7-3.tiff

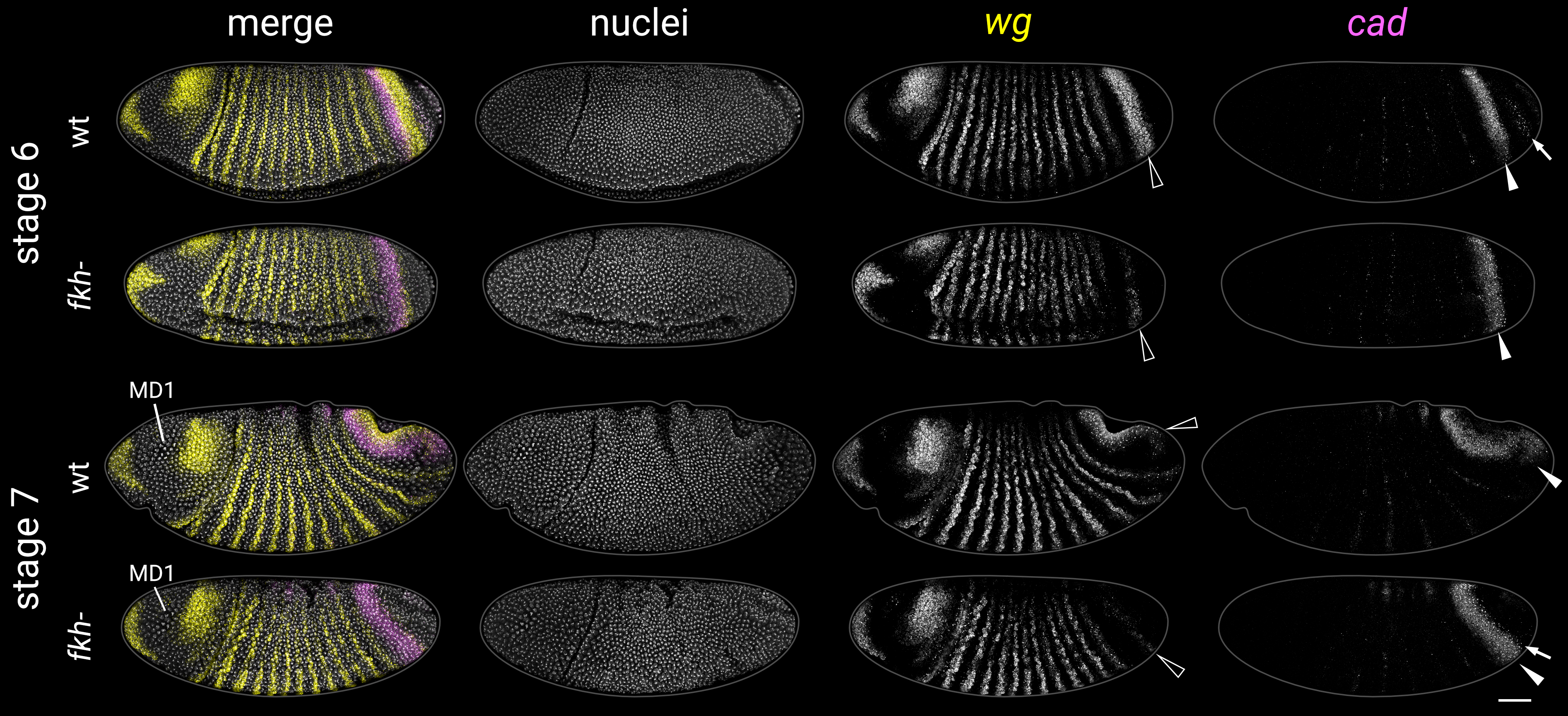

### s7-4.tiff

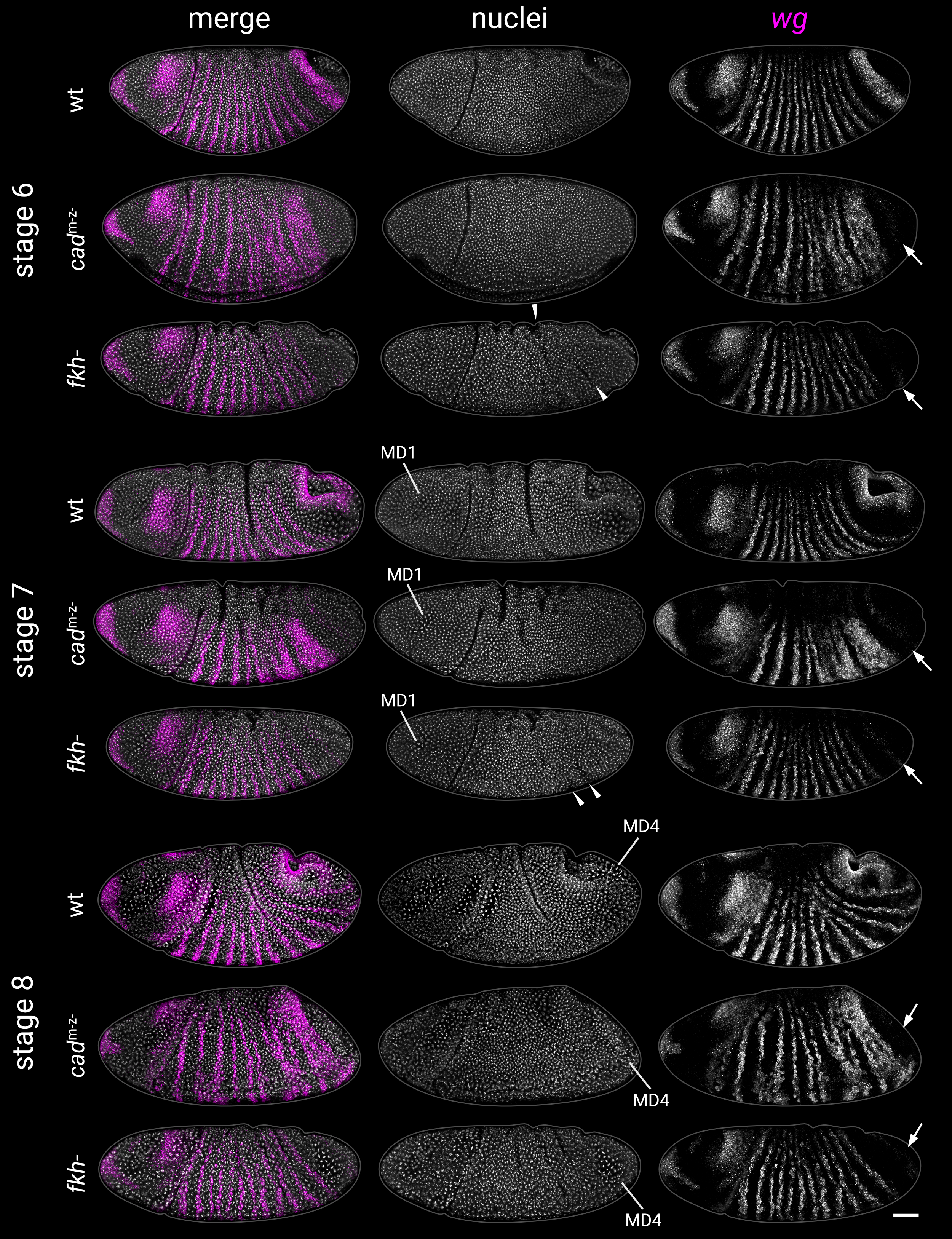

### s8-1.tiff

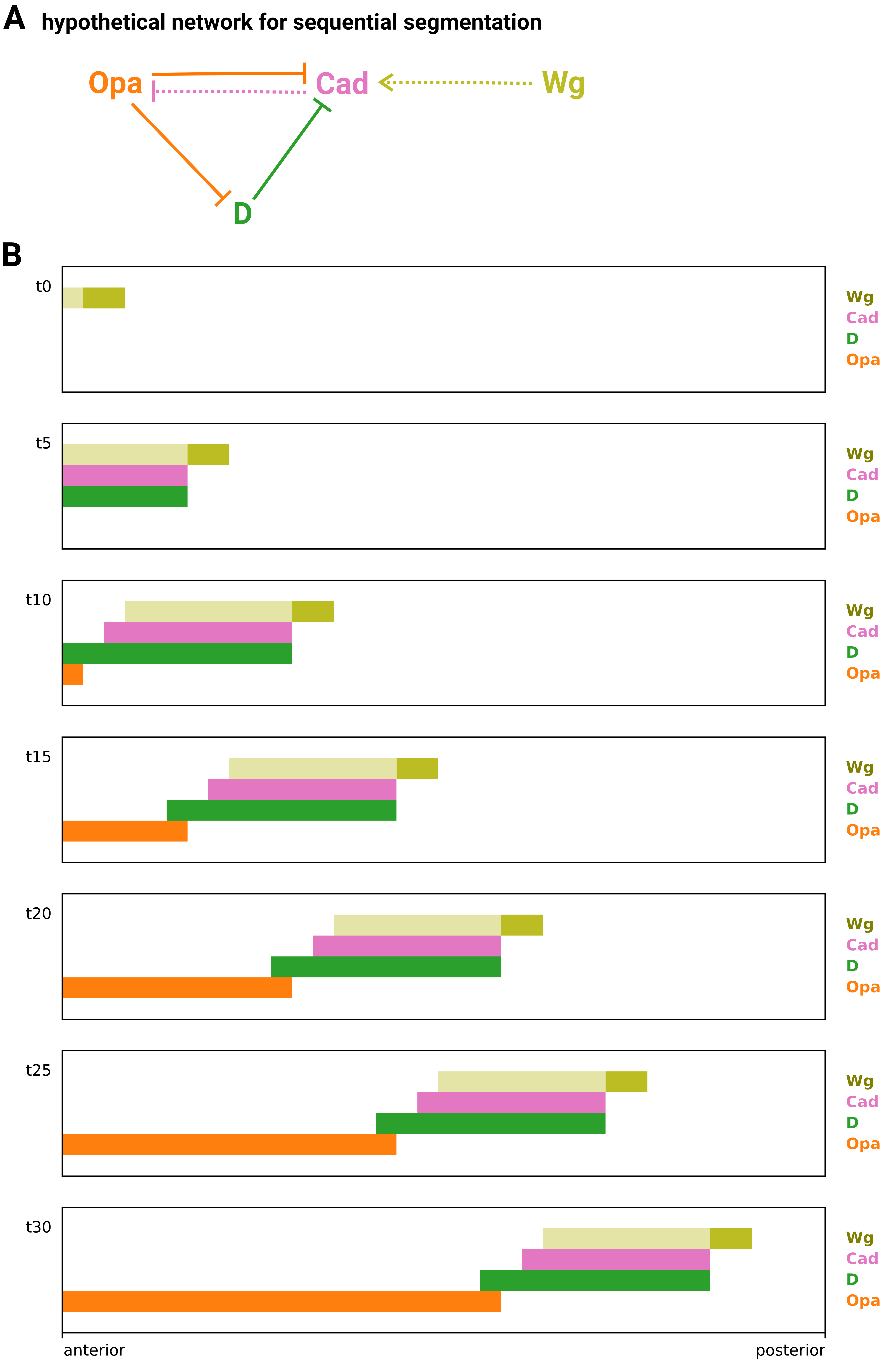
