## Supplementary material for "A timer gene network is spatially regulated by the terminal system in the *Drosophila* embryo": Source Data: fig5-sd2.pdf

**Figure 5—source data 2: width of the posterior *tll* expression domain at stage 5.2 and stage 5.5.** The width of the posterior *tll* domain was measured by counting (manually in confocal $z$-stacks) the number of individual nuclei within a single row of nuclei spanning across the domain on the lateral side of the blastoderm. All embryos were stained in the same tube and imaged in the same session with the same acquisition settings. The width of the domain was greater in the stage 5.2 embryos (mean = 18.7 nuclear diameters) than in the stage 5.5 embryos (mean = 15.0 nuclear diameters).

| stage | embryo | width of *tll* domain (# of nuclei) |
|---|---|---|
| 5.2 | 1 | 19 |
| 5.2 | 2 | 17 |
| 5.2 | 3 | 20 |
| 5.5 | 4 | 16 |
| 5.5 | 5 | 14 |
| 5.5 | 6 | 15 |
